# Toward Textile-Integrated Electrochemical Systems: A Flexible PCB Potentiostat for Wearable Glucose Monitoring

**DOI:** 10.64898/2026.02.24.707657

**Authors:** Sheng Yong, Hassan Hamidi, Daniela Iacopino, Stephen Beeby

**Author notes:** Corresponding authors: S. Y and H. H; email addresses &. S.Y. and H.H. contributed equally to this work.

## Abstract

Flexible and textile-integrated electrochemical systems offer a convenient, user-friendly and non-invasive platform for continuous biochemical monitoring. In this study, a fully flexible and low-profile electrochemical system was developed by fabricating both the glucose biosensor and a compact potentiostat implemented on a polyimide (PI) filament circuit. The glucose biosensor was realized via direct laser writing (DLW), enabling precise electrode patterning and seamless integration with the potentiostat filament circuit. The integrated system exhibited a linear chronoamperometric response to glucose concentrations ranging from 0 to 0.25 mM in artificial sweat (AS). Further evaluation on cotton textiles soaked in AS and under mechanical bending confirmed stable performance, flexibility, and robustness.

These findings highlight the potential of the PI-based potentiostat–sensor system for wearable, textile-integrated glucose monitoring and broader healthcare applications.

## 1 Introduction

Monitoring glucose levels in body fluids is critical for managing diabetes and maintaining overall metabolic health. Conventional glucose measurement techniques typically rely on invasive or semi-invasive sampling methods, which limit their suitability for continuous or user-friendly monitoring.^1,2^ However, in recent years, wearable non-invasive glucose sensing *via* sweat has emerged as a promising alternative, offering a convenient and real-time approach to metabolic analysis.^3,4^ In parallel, in recent years electronic textiles (e-textiles) have become a promising platform for personalized healthcare due to their flexibility, comfort, and ability to support continuous physiological monitoring. E-textiles involve the integration of electronic devices and materials within or on top of a fabric, and can be realized using a range of fabrication techniques, including solution processing and flexible circuit technologies.^5,6^ Numerous wearable textile-based glucose sensor platforms have been recently fabricated using natural (silk and cotton) and polymer-based (polyester, polyurethan) films, threads and fibers demonstrating detection in artificial and real biological fluids in the relevant physiological concentration range. ^7–13^ For example, our group recently demonstrated non enzymatic glucose sensing using a three-yarn electrode system comprising a copper oxide and activated carbon-coated copper working electrode yarn and glassy carbon-coated copper yarns as reference and counter electrodes, achieving a sensitivity of 247.6 μA.^14^ Chowdhury *et al*. developed a textile-based sweat glucose sensor using commercially available conductive Adafruit textiles achieving sensitivity of 0.0019 μA μM^−1^ and an LOD of 63.1 μM.^15^ Zhang *et al*. developed a sweat glucose sensor exploiting the quick-drying quality of Janus fabric, allowing component directly contacting skin to transfer sweat toward the detection site.^16^ The sensor provided accurate glucose level in the testing samples and during human testing during exercise.^16^ Similarly, the incorporation of a hydrophilic textile interface between the sensor working electrode and the skin, has been exploited by various research groups for mitigating enzyme degradation, contamination, and signal drift over time arising from direct contact between the biosensor and the skin in wearable scenarios.^17^

A crucial step for practical applications is the integration of glucose biosensors into e-textiles, along with compact, flexible readout circuits, enabling non-invasive, continuous and real-time glucose monitoring for minimum discomfort diabetes management. Beyond clinical applications, such systems would offer user-friendly platforms for sports and fitness, where real-time glucose tracking can inform energy expenditure and endurance. To date, most reported e-textile glucose sensors rely on external readout systems, such as benchtop potentiostats,^18^ rigid printed circuit boards (PCBs),^19–21^ or transistor-based circuits, to convert,^22^ amplify, and filter the output current signal. These readout systems are often bulky, rigid, and power-demanding, making them unsuitable for seamless textile integration. Transistor-based circuits, in particular, require multiple supply voltages, microcontrollers, or charge pumps, which increase power consumption and design complexity—contradicting the design principles of wearable and energy-efficient electronics. Recently, Huang *et al*. reported a wearable non-invasive glucose monitoring system using a flexible PCB integrated with a metal–organic framework-based sensor patch.^23^ Although the device achieved a sensitivity of 25.48 μA mM□^1^, its circuit design remained complex, comprising over seven rigid chips, and its mechanical flexibility was not rigorously characterized.

In the last ten years Direct Laser Writing (DLW) has emerged as a promising technology for the realization of flexible biosensors. The resulting Laser Induced Graphene (LIG) electrode materials have been shown particularly suited for glucose sensing due to their high electrical conductivity, large surface area, rich surface chemistry and scalable fabrication directly on flexible substrates.^24,25^ LIG can be produced through a one-step laser engraving process on polymeric or textile precursors, enabling the creation of porous, mechanically robust, and highly conductive graphene networks suitable for wearable biosensing applications.^26–28^ In glucose sensing platforms, LIG provides an excellent electrode interface for enzyme immobilization or non-enzymatic catalytic detection, facilitating rapid electron transfer and high sensitivity.^29,30^ When integrated into sensing architectures, LIG-based sensors offer flexibility and mechanical durability, making them highly attractive for continuous, non-invasive monitoring of glucose in sweat. Although These advantages position LIG as a key material for next-generation wearable e-textile biosensors with potential for real-time metabolic health monitoring very few examples of fully integrated LIG-based glucose sensors have been proposed .^31^

In this study, we present a simplified, flexible filament PCB (fPCB) potentiostat circuit for glucose sensing, designed for integration with laser-fabricated glucose sensors. The glucose sensor was fabricated using direct laser writing (DLW) on flexible polyimide (PI) tapes, enabling precise electrode patterning with minimal processing steps. The circuit employed only two compact Mini Small Outline Package (MSOP)-8 chips (3 mm × 3 mm × 0.65 mm) functioning as the biosensor control element and transimpedance amplifier, complemented by a few passive components. Operating from a single 3.7 V coin cell, the design eliminates the need for charge pumps, negative supply voltages, or additional power-conditioning circuits. The resulting system displayed low power consumption, high mechanical flexibility, and a compact form factor, utilizing a 50 μm thick PI substrate with 35 μm thick copper tracks. The platform was evaluated for glucose detection in artificial sweat (AS) and further tested when integrated onto cotton textiles soaked in AS and under mechanical bending. Results confirm that the fully flexible potentiostat–sensor system maintained stable electrochemical performance and mechanical resilience, demonstrating its potential as a wearable, textile-integrated platform for continuous glucose monitoring in sustainable healthcare applications.

## 2 Experimental section

### 2.1 Materials

Glucose oxidase (GOx) type VII (EC 1.1.3.4, Aspergillus niger, 221 U mg-1), D(+)-glucose, bovine serum albumin (BSA), chitosan (CS, medium molecular weight), Nafion perfluorinated resin (5%W/W), uric acid, urea, lactic acid, ascorbic acid, sodium chloride, ammonium chloride, tetrathiafulvalene (TTF), and Ag|AgCl paste were procured from Sigma-Aldrich. Polyimide tape (3M, thickness ca. 70 μm) was purchased from Radionics.

### 2.2 Characterization

The morphology of the LIG surface was investigated via scanning electron microscopy (Zeiss 16 SUPRA 35 VP operating at 5kV). The LIG Raman characteristics were investigated using a Horiba XploRA Raman microscope equipped with a 70 mW, 532 nm laser.

### 2.3 Glucose biosensor strips fabrication

The glucose biosensor strips were fabricated using direct laser writing (DLW) following previously reported procedures with minor modifications.^18^ Laser-induced graphene (LIG) electrodes were fabricated using a handheld laser engraver (LaserPecker2, 5 W, 450 nm) operated at 2k resolution, 14% laser power, 14% engraving depth, and a 5 mm defocusing distance. After laser scribing, residual carbon debris was removed by sequential rinsing with acetone, isopropyl alcohol, and deionized water. A three-electrode configuration (23 mm × 6 mm) comprising a 2 mm diameter circular working electrode (WE), a counter electrode (CE), and a reference electrode (RE) was patterned onto PI tape. The width of the PI tape was reduced to 12 mm to improve compatibility with textile manufacturing and potential weaving integration. The RE was prepared by coating the designed LIG electrode area with Ag/AgCl paste, followed by curing at 100 °C for 10 min. The WE was modified with tetrathiafulvalene (TTF) by drop-casting 2 μL of a 15 mg mL□^1^ TTF solution in tetrahydrofuran (THF) and allowing the solvent to evaporate at room temperature. For enzyme immobilization, glucose oxidase (GOx, 5 U μL□^1^ in phosphate-buffered saline, PBS), bovine serum albumin (BSA, 10 mg mL□^1^ in PBS), and chitosan (1% w/v in 1% v/v acetic acid) were mixed in equal volumes. A 10 μL aliquot of the mixture was drop-cast onto the LIG/TTF-modified WE and dried at room temperature. Subsequently, 5 μL of Nafion solution was applied as a protective coating layer. The completed electrodes were dried and stored under refrigerated conditions until use. These devices are hereafter referred to as glucose biosensor strips.

### 2.4 fPCB design/fabrication

The circuit layout was designed using EAGLE 9.6.2 (Autodesk, USA), and the flexible printed circuit board (fPCB) was fabricated by JLCPCB, Inc. (China). The potentiostat circuit employed a dual operational amplifier (LMP2232) for control and voltage follower functions, and a transimpedance amplifier (OPA380) to convert the current generated between the working and reference electrodes into a voltage signal. In the initial V1 fPCB, TL072 operational amplifiers were used for control, voltage follower functions, and a transimpedance amplifier. The transimpedance amplifier was configured with a gain of 100 kΩ and a low-pass filter with a cutoff frequency of 2 Hz. All components were powered using a single 3.7 V supply. The output signal was recorded using an oscilloscope and analyzed using OriginPro 2023 (OriginLab, USA).

### 2.5 Measurement

Chronoamperometry (CA) measurements of the biosensor strips were performed using a commercial benchtop potentiostat (PGSTAT101, Metrohm Autolab, The Netherlands). Phosphate buffered saline (PBS, pH 7.0) and artificial sweat (AS) were used as electrolyte solutions. Electrochemical measurements using the fPCB potentiostat and the biosensor strips were conducted with a 3.7 V DC power supply and an applied bias voltage of 0.4 V generated using a resistive voltage divider. The output signals were recorded using a digital storage oscilloscope (DSO1002A, Agilent Technologies) with an input resistance of 1 MΩ. For textile integration experiments, the sensor surface was covered with a 10 mm × 10 mm cotton textile sample pre-soaked with the test solution. Signal acquisition during textile testing was performed using a data logger (PicoLog 1216, Pico Technology). Mechanical bending tests were conducted by folding the fPCB potentiostat while performing measurements in artificial sweat containing glucose.

## 3 Results and discussion

Figure 1a illustrates the fabrication process of the LIG electrodes via DLW on thin PI tape, followed by sequential drop-casting of TTF, GOx and Nafion onto the LIG WE to form the glucose biosensor. Due to the miniaturization of features, the morphology of the scribed LIG material was characterized using SEM prior surface modification to ensure that the morphology of the resulting LIG electrodes was suitable for further biosensor fabrication. The SEM image of Figure 1b revealed the characteristic porous LIG structure composed of a dense interconnected three-dimensional carbon network, showing morphological features like other reported LIG structures ^18,32^. This highly porous architecture is crucial to provide high electroactive surface area and to promotes efficient electron transfer, both essential for sensitive electrochemical detection. The LIG morphology was further characterized by Raman spectroscopy. Figure 1c showed that the LIG spectrum exhibited the characteristic features of graphitic materials, with D, G, and 2D peaks centered at 1345, 1580, and 2689 cm^-1^, respectively, corresponding to defects in graphene, sp^2^ carbon stretching, and second-order zone-boundary phonons.^33^ The I_D_/I_G_ ratio of 0.95 (±0.5) was associated with high disorder and heterogeneity, while the I_2D_/I_G_ ratio of 0.85 (±0.2) indicated the presence of multilayer graphene structures.^34,35^ A photograph of the resulting flexible three-electrode biosensor is shown in the inset of Figure 1c. The analytical performance of the miniaturized sensing platform was first evaluated using chronoamperometry (CA) measurements performed with a commercial potentiostat across glucose concentrations ranging from 0.05 to 0.6 mM in PBS buffer. This concentration range encompasses physiologically relevant glucose levels typically reported in human sweat (0.05–0.2 mM). As shown in Figure 1d, the biosensor current response increased systematically with increasing glucose concentration, confirming effective enzymatic glucose oxidation and signal transduction. The corresponding calibration plot (Figure 1d, inset) exhibited a strong linear relationship between current response and glucose concentration, yielding a sensitivity of 36.5 ± 1.5 μA mM□^1^ cm□^2^ with a correlation coefficient (R^2^) of 0.994. The limit of detection (LOD, S/N = 3) was calculated to be 0.033 mM. These analytical metrics are comparable to those previously reported for larger-format LIG glucose sensors on PI, indicating that device miniaturization did not compromise electrochemical performance. This result highlights the suitability of the platform for wearable and textile-integrated sensing applications.

**Figure 1.**
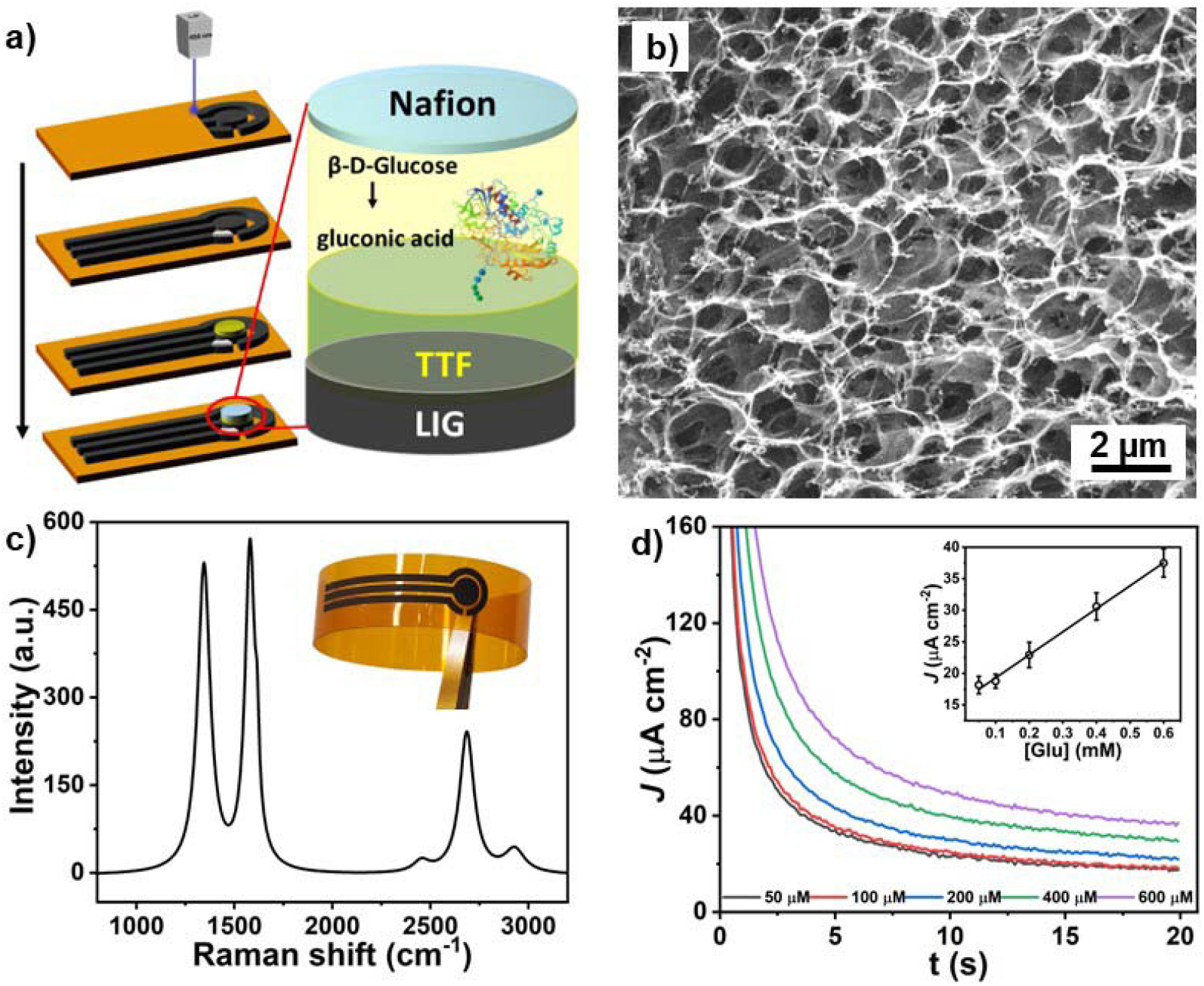
a) Schematic of DLW fabrication of the three-electrode feature on PI following by biofunctionalization steps with details of biosensor architecture and glucose oxidation mechanism; b) SEM image of LIG structure; c) Representative Raman spectrum of LIG. Inset: photograph of LIG electrodes on flexible PI; d) Plot of CA response of LIG biosensor at different glucose concentrations (50 - 600 mM) in PBS solution (pH 7) at an applied potential of +0.25 V. Inset: corresponding calibration curve.

The custom potentiostat was developed through four design generations (V1–V4), with each iteration addressing key challenges related to circuit performance, miniaturization, and mechanical flexibility. All versions were implemented on PI-based flexible substrates to maintain compatibility with wearable and textile-integrated sensing platforms. Photographs of all the designs are shown in Figure 2 (a-d), with detailed circuit schematics and specifications provided in the Supplementary Information. The initial V1 fPCB (see Figure 2a) served as proof-of-concept device and demonstrated reliable potentiostat functionality and signal conversion. However, this configuration (TL072 chip) required dual ±5 V supplies increasing power consumption and circuit complexity, which limited its suitability for wearable applications. Building upon this foundation, the subsequent V2 and V3 designs incorporated integrated amplifier components (LMP2322) capable of operating under a single +3.7 V supply, significantly simplifying circuit complexity and improving mechanical compliance. The V2 design (Figure 2b), incorporated a dedicated transimpedance amplifier (OPA 380) together with control and voltage follower stages, as well as passive low-pass filtering to improve signal stability. A passive low-pass filter was also added at the output to improve signal quality. This design simplified power management, however, the relatively large component footprint (SOIC-8 package (4 mm × 6.2 mm × 1.35 mm) of the OPA380, along with surface-mounted capacitors (0603 package) at the output), restricted the miniaturization potential for flexible strip applications. The improved V3 design (Figure 2c) addressed this limitation by adopting smaller MSOP-packaged components (3 mm × 3 mm × 0.65 mm) and introducing a connector interface to enable direct attachment of the biosensor strip. These optimizations significantly reduced the width of fPCB strip from 25 mm to 15 mm. However, the connector’s geometry imposed constraints that limited further dimensional reduction. The final V4 potentiostat represents the fully optimized version, achieving a compact and narrow form factor (6 mm × 40 mm) suitable for integration into textile substrates (Figure 2d). Compared with earlier iterations, the V4 design demonstrated enhanced mechanical flexibility, reduced power consumption, and improved operational stability. These characteristics support reliable electrochemical signal acquisition in wearable and textile-based glucose sensing applications

**Figure 2.**
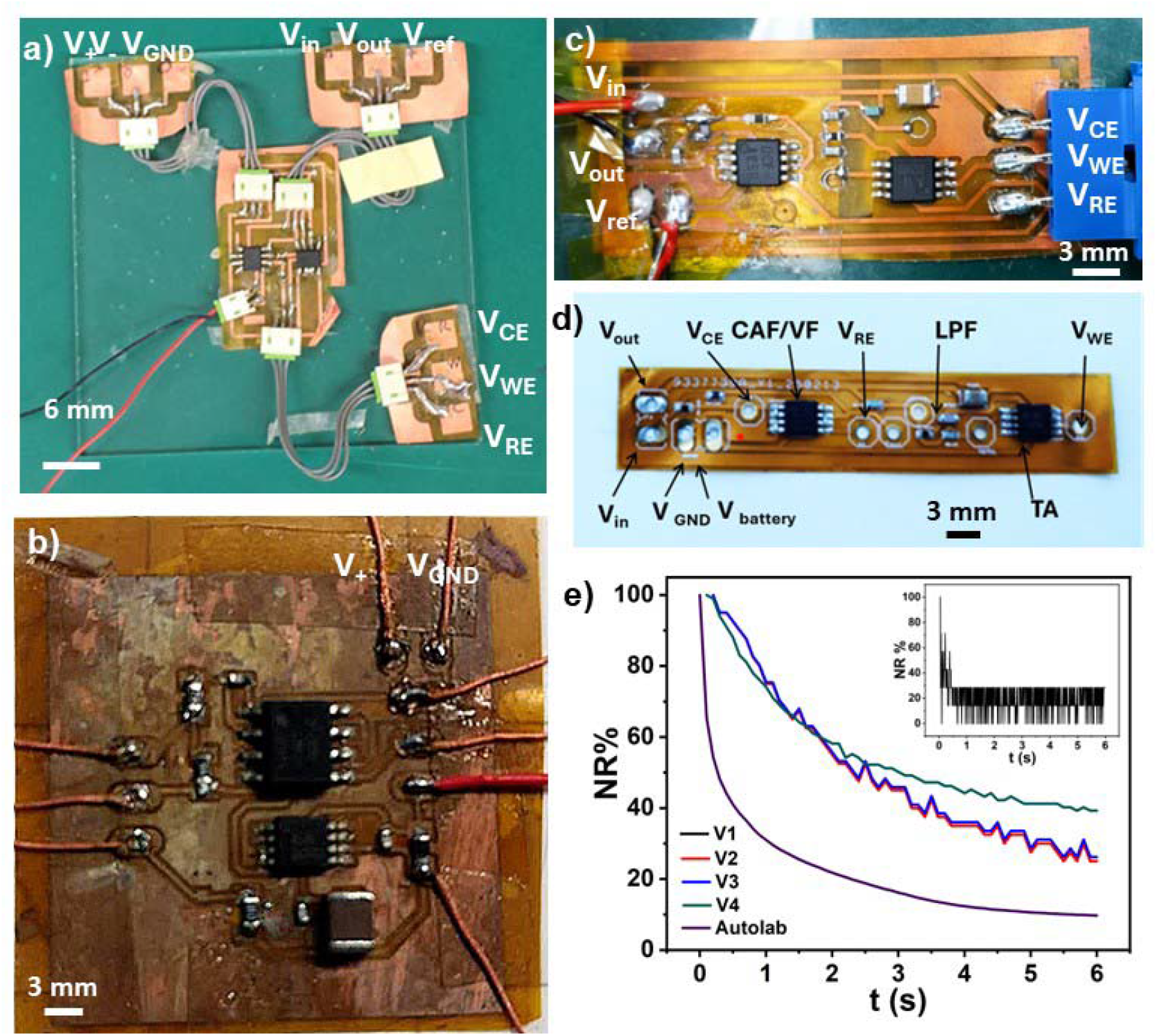
Photographs of (a) V1, (b) V2, (c) V3, and (d) V4 fPCBs showing progressive miniaturization and simplification. (e) Normalized chronoamperometric responses to 1 mM glucose recorded in PBS at an applied potential of +0.25 V using fPCBs V1–V4 and compared with the benchtop Autolab response.

The normalized chronoamperometric responses obtained from the different fPCB potentiostat generations (V1–V4) were compared with those recorded using a commercial benchtop potentiostat, as shown in Figure 2e. Measurements were performed using 1 mM glucose in PBS. The responses were normalized to their initial current values to account for differences in input impedance between the benchtop potentiostat (with an input resistance of 100 GΩ) and the digital storage oscilloscope used for fPCB signal acquisition (with an input resistance of 1 MΩ). All fPCB configurations exhibited similar current response trends. However, the V1 design (Figure 2d inset) showed significant signal noise, attributed to instability in the dual power supply configuration and parasitic capacitance effects. The V2 and V3 potentiostat versions demonstrated improved signal quality but still exhibited higher noise levels compared to later designs, likely due to suboptimal circuit layout and component value selection. Following the initial capacitive discharge phase (∼1 s), the chronoamperometric response obtained from the V4 FPCB closely paralleled that of the commercial potentiostat. This agreement confirms the reliability, stability, and analytical performance of the optimized flexible potentiostat, supporting its suitability for portable and textile-integrated electrochemical glucose sensing applications.

Following optimization through successive design iterations, the final flexible potentiostat (V4) was selected for electrochemical performance evaluation under physiologically relevant conditions. The electronic readout system (Figure 3a,b) consists of a multistage signal conditioning architecture designed for electrochemical biosensing. The input stage incorporates analog amplification circuitry that processes the low-level sensor current generated by the electrochemical reaction, providing initial signal amplification and stabilization. The amplified signal is then routed to a potentiostatic control block, which maintains a constant potential between the WE, RE and CE. This module regulates the electrode bias through feedback control, ensuring stable electrochemical operation and accurate current measurement. The following stage includes an additional amplification and offset regulation unit powered by a battery supply, which further increases signal magnitude while maintaining proper grounding and noise suppression. Finally, the conditioned analog output signal is passed through a low-pass filter to remove high-frequency noise and interference, producing a stable output suitable for data acquisition and subsequent signal processing. The weight of the fully integrated e-textile sensor is only 0.7 g. The cost of the V4 fPCB is approximately $15 (OpA 380∼$7.5 LMP2322 $2, wire + PCB $5 and $0.5 for 402 resistor and caps Relative to a standard over-the-counter glucose meter (≈$50), our integrated system costs only ≈50%, underscoring its exceptional affordability.

**Figure 3.**
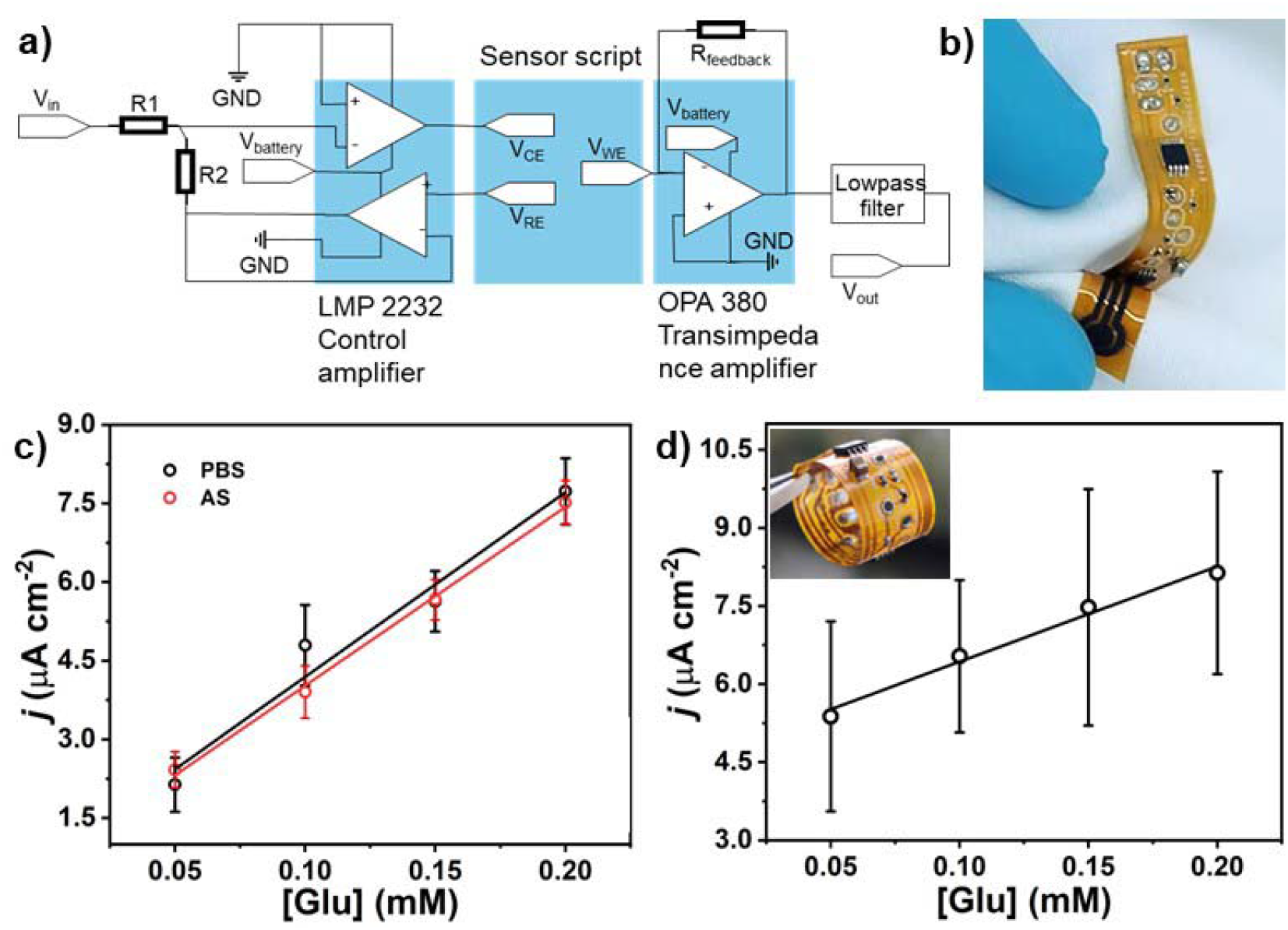
a) System-level block diagram of the fPCB; b) Photograph of the fully integrated biosensor using V4-fPCB connected with the glucose biosensor strip; c) Calibration plots for glucose obtained in PBS and AS solution at increasing glucose concentrations (0-0.2 mM); d) Calibration plot in AS solution obtained using integrated biosensor with folded fPCB. Inset; photograph of the bent fPCB.

To assess its suitability for wearable sensing applications, the capabilities of the integrated glucose biosensor strip were evaluated in buffer (PBS) and artificial sweat (AS). Figure 3c shows calibration plots obtained from chronoamperometric measurements performed in both PBS and AS containing glucose concentrations ranging from 0 to 0.20 mM. In both media, the measured current density increased proportionally with glucose concentration, indicating stable and reproducible electrochemical responses. The calibration plots exhibited strong linear relationships, with sensitivities of 35.2 ± 4.7 μA mM^-1^ cm^-2^ in PBS and 34.10 ± 1.2 μA mM^-1^ cm^-2^ in artificial sweat, and correlation coefficient (R^2^) values of 0.966 and 0.998, respectively. These analytical performances were comparable to those obtained using a benchtop potentiostat, demonstrating that miniaturization and flexibility did not compromise sensing accuracy or stability. Statistical comparison of the calibration slopes at the 0.05 significance level revealed no significant difference between responses obtained in PBS and artificial sweat, confirming equivalent sensor sensitivity in both media and consistent with prior reports ^12^.

The mechanical robustness of the integrated system was further evaluated under folded conditions (Figure 3c). While mechanical deformation of the fPCB resulted in a reduced signal amplitude and a lower sensitivity of 18.3 ± 1.6 μA mM□^1^ cm□^2^, the calibration response remained highly linear (R^2^ = 0.984). This behavior indicates that, despite mechanical strain, the device preserved proportional glucose detection and functional electrochemical performance. Collectively, these results validate the potential of the integrated flexible glucose biosensor strip for wearable and textile-integrated glucose monitoring in continuous and sustainable healthcare applications.

Building on the successful electrochemical validation of the flexible biosensor in AS, the system was further evaluated for textile integration to simulate realistic wearable conditions.

Figure 4 presents the performance of the sensing platform when glucose-loaded cotton textiles were placed directly onto the biosensor strip integrated with the fPCB. As illustrated in Figure 4 inset, cotton samples were soaked in AS containing defined glucose concentrations (0-0.2 mM) and subsequently positioned over the sensing region for electrochemical measurements. The resulting calibration curves showed a clear linear increase in current density with increasing glucose concentration for both artificial sweat alone (AS-fPCB) and cotton soaked in artificial sweat (AS-fPCB_cotton). Linear regression analysis yielded sensitivities of 32.9 ± 5.4 μA cm□^2^ mM□^1^ for AS-FPCB and 32.6 ± 4.3 μA cm□^2^ mM□^1^ for AS-FPCB_cotton. The closely overlapping current responses and comparable calibration slopes indicate that the textile interface did not significantly affect electrochemical signal transduction or sensor sensitivity. These findings demonstrate that the fully flexible electrochemical sensing system can be effectively integrated into textile materials while maintaining high analytical performance, mechanical resilience, and signal stability. This capability represents an important step toward the development of fully wearable, textile-based glucose sensing platforms for continuous and sustainable health monitoring.

**Figure 4.**
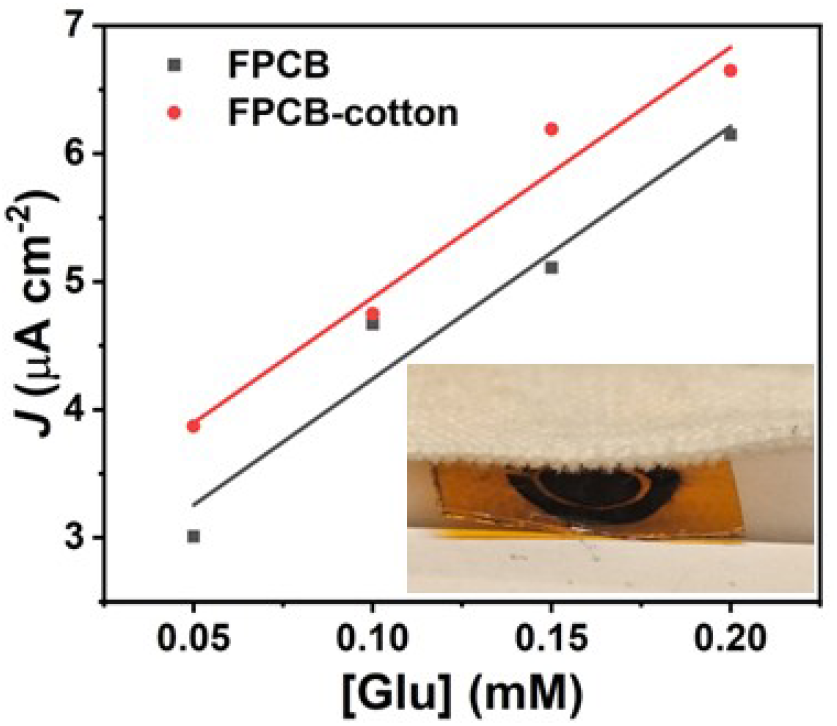
Plot of current density versus glucose concentration of the integrated system in AS solution (black) and cotton (red, soaked with AS) collected using Pico log datalogger. Inset: photograph of integrated biosensor with glucose-loaded cotton textile.

## 4 Conclusions

A narrow, fully flexible PI platform integrating a LIG glucose biosensor with a compact flexible PCB potentiostat was developed for non-invasive glucose monitoring. The integrated system exhibited stable and linear electrochemical responses in both PBS and AS, with analytical performance closely matching that of a commercial benchtop potentiostat. When evaluated on cotton textiles soaked in AS and under mechanical bending, the platform maintained consistent performance, confirming its mechanical robustness, flexibility, and operation reliability. This work demonstrates a scalable and versatile design strategy for realizing self-contained, textile-compatible electrochemical sensing systems. The proposed platform can be readily extended to other biosensing analytes and wearable health-monitoring applications, representing a significant step toward next-generation sustainable, flexible, and fully integrated e-textile platforms. Future developments will focus on optimizing sweat management layers, improving long-term mechanical durability, and integrating wireless data transmission and power management modules. Collectively, these advancements will support the realization of fully autonomous, textile-based glucose monitoring systems for continuous and non-invasive health monitoring.

## 5 Acknowledgement

This publication has emanated from research conducted with the financial support of the Royal Academy of Engineering under the Chairs in Emerging Technologies Scheme, UK Research and Innovation (UKRI) through the EU Horizon Guarantee Scheme for Sustainable Textile Electronics (STELEC), and the European Union’s Horizon Europe programme under the projects INFRACHIP (Grant No. 101131822), GreenArt (Grant No. 101060941), and Herit4Ages (Grant No. 101123175). Additional support was provided by Science Foundation Ireland through the European Regional Development Fund (Grant No. 13/RC/2077-P2, CONNECT) and by the European Union’s Horizon 2020 programme under the Marie Skłodowska-Curie Actions (SusBioLIG, Grant No. 101032167).

